# Epigenetic signatures of starting and stopping smoking

**DOI:** 10.1101/402453

**Authors:** Daniel L McCartney, Anna J Stevenson, Robert F Hillary, Rosie M Walker, Mairead L Bermingham, Stewart W Morris, Toni-Kim Clarke, Archie Campbell, Alison D Murray, Heather C Whalley, David J Porteous, Peter M Visscher, Andrew M McIntosh, Kathryn L Evans, Ian J Deary, Riccardo E Marioni

## Abstract

**Background:** Multiple studies have made robust associations between differential DNA methylation and exposure to cigarette smoke. But whether a DNA methylation phenotype is established immediately upon exposure, or only after prolonged exposure is less well-established. Here, we assess DNA methylation patterns in current smokers in response to dose and duration of exposure, along with the effects of smoking cessation on DNA methylation in former smokers.

**Methods:** Dimensionality reduction was applied to DNA methylation data at 90 previously identified smoking-associated CpG sites for over 4,900 individuals in the Generation Scotland cohort. K-means clustering was performed to identify clusters associated with current and never smoker status based on these methylation patterns. Cluster assignments were assessed with respect to duration of exposure in current smokers (years as a smoker), time since smoking cessation in former smokers (years), and dose (cigarettes per day).

**Results:** Two clusters were specified, corresponding to never smokers (97.5% of whom were assigned to Cluster 1) and current smokers (81.1% of whom were assigned to Cluster 2). The exposure time point from which >50% of current smokers were assigned to the *smoker-enriched* cluster varied between 5-9 years in heavier smokers and between 15-19 years in lighter smokers. Low-dose former smokers were more likely to be assigned to the *never smoker-enriched* cluster from the first year following cessation. In contrast, a period of at least two years was required before the majority of former high-dose smokers were assigned to the never smoker-enriched cluster.

**Conclusions:** Our findings suggest that smoking-associated DNA methylation changes are a result of prolonged exposure to cigarette smoke, and can be reversed following cessation. The length of time in which these signatures are established and recovered is dose dependent. Should DNA methylation-based signatures of smoking status be predictive of smoking-related health outcomes, our findings may provide an additional criterion on which to stratify risk.

## Background

Cigarette smoking is among the leading causes of illness and premature death worldwide [1]. In addition to multiple cancers [2], it is a major risk factor for cardiovascular and respiratory disorders [3, 4]. Recent studies suggest that altered DNA methylation may play an important role in the biological pathways linking smoking to adverse health outcomes [5, 6, 7, 8, 9].

DNA methylation is an epigenetic modification, typically characterised by the addition of a methyl group to a cytosine-guanine dinucleotide (CpG). Both genetic and environmental factors can modulate DNA methylation levels, which in turn can regulate gene expression [10]. To date, the most informative environmental correlate of DNA methylation has been cigarette smoking. Multiple epigenome-wide association studies (EWAS) have been performed on smoking, using either status (e.g. current smoker, former smoker, never smoker) or intake (e.g. pack years) as the trait of interest [5, 7, 9], identifying thousands of smoking-associated loci. Moreover, cohort studies have reported altered DNA methylation in the offspring of women who smoked during pregnancy [11, **Error! Reference source not found.**, 13]. These analyses have identified a large number of loci where methylation is altered by exposure to cigarette smoke, with the cg05572921 locus in the aryl hydrocarbon receptor (AHR) repressor (*AHRR*) gene being among the most robustly implicated [6, 7, 8, 13]. The relationship between exposure to cigarette smoke and DNA methylation changes has been widely reported. However, when these effects are established in smokers and whether they can be recovered by cessation is not well understood. Studying the mechanics of smoking-associated DNA methylation changes may provide a novel means of identifying risk of smoking-related morbidities.

We investigated the extent to which smoking-associated DNA methylation changes were associated with duration of exposure in current smokers and time since cessation in former smokers. We examined the relationship between DNA methylation and smoking in a cohort of over 4,900 individuals, incorporating self-reported years as a smoker and cigarettes per day as metrics for duration of exposure and dose, respectively.

## Methods

### The Generation Scotland Cohort

Details of the Generation Scotland: Scottish Family Health Study (GS:SFHS) have been described previously [14, 15]. DNA samples were collected for genotype- and DNA methylation-profiling along with detailed clinical, lifestyle, and sociodemographic data. The current study comprised 4,929 individuals from the cohort for whom both DNA methylation and smoking data were available. A summary of variables assessed in this analysis is presented in **Table 1**.

**Table 1:** **Summary of the Generation Scotlan d cohort and variable s assessed**

| **Variable** | **N** | **Mean** | **SD** |
| --- | --- | --- | --- |
| **Sex** |  |  |  |
| Males | 1,872 | - | - |
| Females | 3,033 | - | - |
| Age (years) | 4,905 | 48.5 | 13.9 |
| **Smoking status** |  |  |  |
| Current smoker | 917 | - | - |
| Former smoker | 1,466 | - | - |
| Never smoker | 2,522 | - | - |
| **Smoking variables** |  |  |  |
| Cigarettes per day (current and former smokers) | 2,177 | 11.1 | 9.8 |
| Cigarettes per day (current smokers) | 859 | 15.2 | 9.8 |
| Cigarettes per day (former smokers) | 1,318 | 8.5 | 8.8 |
| Pack years (current and former smokers) | 2,037 | 15.9 | 16.8 |
| Pack years (current smokers) | 854 | 23.3 | 19.7 |
| Pack year (former smokers) | 1,183 | 10.6 | 11.8 |
| Years as a smoker (current and former smokers) | 2,221 | 27.7 | 13.4 |
| Years as a smoker (current smokers) | 917 | 28.7 | 13.2 |
| Years as a smoker (former smokers) | 1,304 | 27.0 | 13.5 |
| Years since cessation (former smokers only) | 1,324 | 9.0 | 10.0 |

**Summary of the Generation Scotland and cohort and variables assessed**

GS:SFHS smoking data were collected using two different questionnaires. The first version of the questionnaire was answered by 2,162/4,929 (43.8%) of the participants and collected data on absolute values with respect to number of cigarettes smoked, and age started/stopped smoking. The second version of the questionnaire, which was answered by the remaining 2,779 individuals in the analysis sample (56.2%), collected data using binned intervals. In order to harmonise the two sets of measurements, mid-point interval estimates were calculated for the ordinal data from the second version of the questionnaire (e.g., 17 years of exposure was assigned to individuals who had reported smoking between 15-19 years). Second-hand smoking status was assigned based on whether participants reported exposure to cigarette smoke at home, work or elsewhere, or whether they reported cohabiting with a smoker. Both questionnaires can be accessed from the GS:SHFS website (www.generationscotland.co.uk).

In the current study, exposure data were placed into ten five-year bins from 0-4 years to 45-49 years (N ≥ 32 per bin), with the longest exposure defined as ≥ 50 years (N = 23). Data on time since cessation were placed into five-year bins from 10-14 years to 30-34 years (N ≥ 48 per bin). The longest time since cessation was defined as ≥35 years (N = 53), whereas the most recent cessation time points (0-9 years) were presented as yearly intervals (N ≥ 26). Sample counts at each exposure and cessation time point are presented in **Supplementary Tables 1-2**.

### Ethics

All components of GS:SFHS received ethical approval from the NHS Tayside Committee on Medical Research Ethics (REC Reference Number: 05/S1401/89). GS:SFHS has also been granted Research Tissue Bank status by the Tayside Committee on Medical Research Ethics (REC Reference Number: 10/S1402/20), providing generic ethical approval for a wide range of uses within medical research.

### GS:SFHS DNA methylation

Genome-wide DNA methylation was profiled in 5,200 individuals using the Illumina HumanMethylationEPIC BeadChip. Quality control was conducted in R [16]. ShinyMethyl [17] was used to plot the log median intensity of methylated versus unmethylated signal per array, with outliers excluded upon visual inspection. WateRmelon [18] was used to remove (1) samples where ≥1% of CpGs had a detection p-value in excess of 0.05 (2) probes with a beadcount of less than 3 in more than 5 samples, and (3) probes where ≥0.5% of samples had a detection p-value in excess of 0.05. ShinyMethyl was used to exclude samples where predicted sex did not match recorded sex. Ten saliva-derived samples and three samples from individuals who had answered “yes” for all self-reported conditions were also excluded. This left a sample of 5,088 blood-derived samples available for analysis, of whom 4,929 had smoking data available.

### Statistical analysis

#### All analyses were performed in R [16]

Data-driven cluster analysis was performed on the first two components, identified via data reduction analysis [19] on the top 100 p-value-ranked methylation sites from a recent, large meta-analysis EWAS of current versus never smoking (Joehanes et al. Supplementary Table 1, Sheet 02) [5]. Ninety of the top 100 probes were present in the GS:SHFS DNA methylation dataset following quality control. Principal coordinates analysis [19] was performed on the data using cmdscale() function in the *stats* package [16]. K-means clustering was performed to partition the data, using the kmeans() function in the *stats* package [16]. As the probe set under consideration was associated with current/never smoker status, two clusters were specified.

Logistic regression was performed to assess the relationship between a genetic variant in the *CHRNA5-A3-B4* gene cluster that is associated with heaviness of smoking (rs1051730) [21] and cluster assignment in current smokers, adjusting for sex. The relationships between cluster assignment and batch, sex, and passive smoking were assessed using Chi-Squared Tests. The relationship between cluster assignment and alcohol consumption (current, former and never drinker) was assessed using a Fisher’s Exact Test.

Data were visualised using “broken stick” regression lines using the default parameters for the segmented() function in the *segmented* package in R [20].

## Results

Descriptive data for the 917 current-, 1,466 ex-, and 2,522 never-smokers are summarised in

### Clustering of current smokers depends on dose and duration smoked

Of the 2,522 never smokers, 2,459 (97.5%) were assigned to a *never smoker-enriched* cluster whereas, of the 917 current smokers, 744 (81.1%) were assigned to a *smoker-enriched* cluster (**Figure 1**). There was no association between misclassification of current smokers to the *never smoker-enriched* cluster and sex, alcohol consumption, batch or genotype at the well-established nicotine addiction genetic variant rs1051730 [21], (P ≥ 0.103; **Supplementary Table 3**). Similarly, there was no association between misclassification of never smokers (n=63) to the *smoker-enriched* cluster and exposure to second-hand smoke, sex, alcohol consumption, or plate processing batch (P ≥ 0.179 **Supplementary Table 3**). The proportion of individuals assigned to the *smoker-enriched* cluster increased with years as a smoker (**Figure 2, Supplementary Table 1**). Of the 32 individuals who reported smoking for 0-4 years prior to DNA methylation sampling, seven (21.9%) were assigned to the *smoker-enriched* cluster; for the 76 individuals who reported smoking for 5-9 years prior to sampling, 34 (44.7%) were assigned to the *smoker-enriched* cluster. The proportion of assignments to the *smoker-enriched* cluster increased to 87.3% for current smokers at 20-24 years of exposure, remaining stable thereafter. Of the 670 current smokers reporting at least 20 years of exposure 605 (90.3%) were assigned to the *smoker-enriched* cluster.

**Figure 1:**
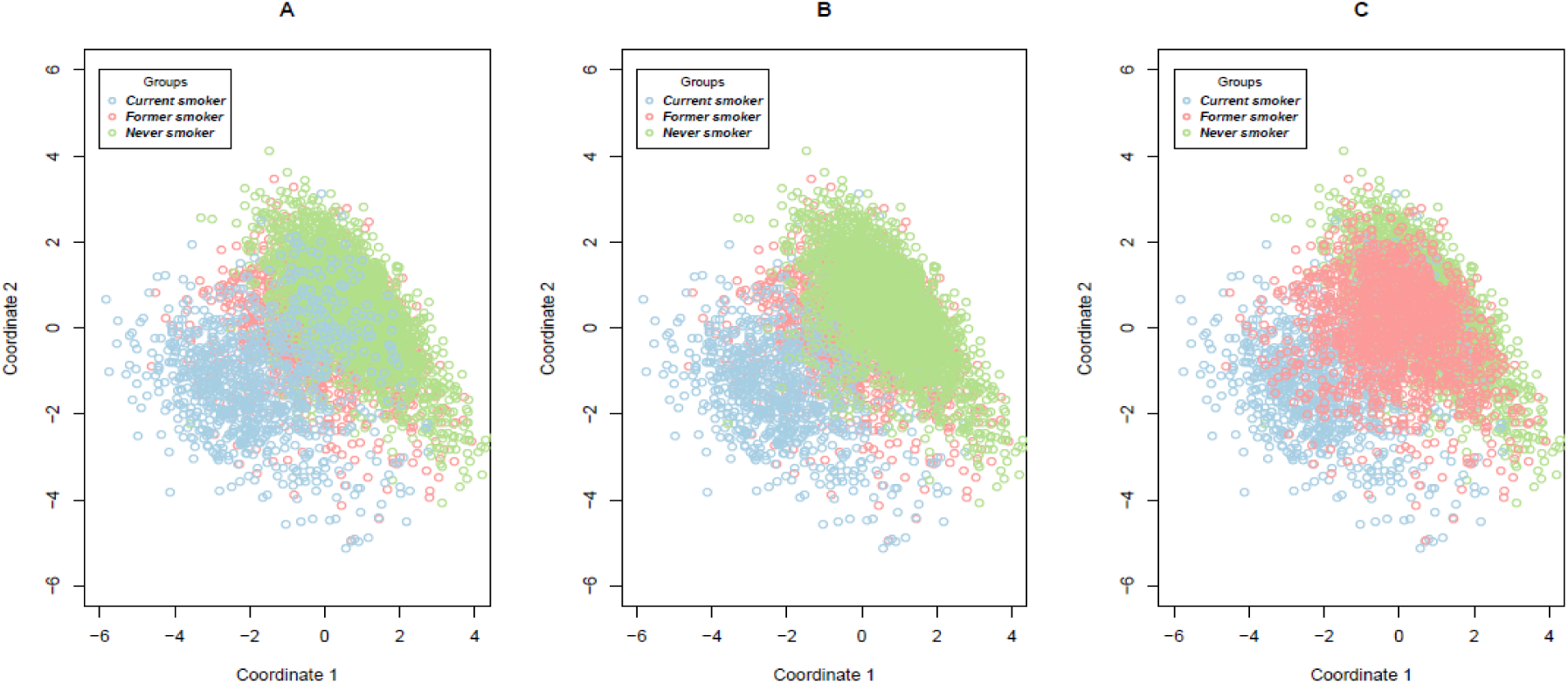
Principal coordinate vectors 1 and 2 from a multidimensional scaling analysis of 90 smoking-associated probes. Samples are coloured by smoking status (blue = current smokers, green= never smokers, pink = former smokers). Panel A presents current smokers in the foreground. Panel B presents never smokers in the foreground, and panel C presents former smokers in the foreground.

**Figure 2:**
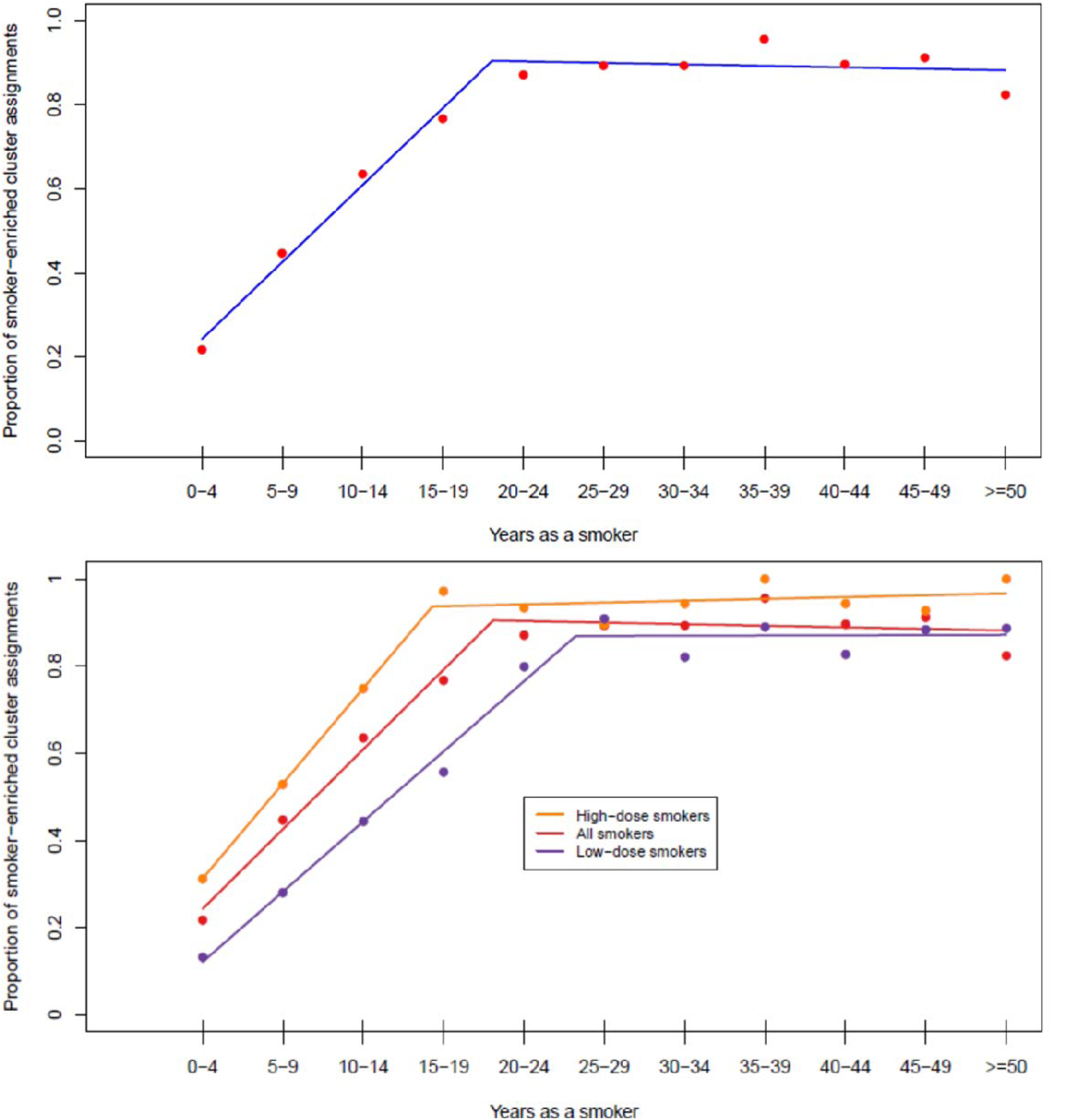
Proportion of current smokers assigned to Cluster 2 (smoker-enriched cluster) by duration of exposure. The upper panel displays a “broken stick” regression line for all samples. The lower panel displays “broken stick” regression lines for all samples (red), high-dose samples (orange) and low-dose samples (purple).

There was a significant association between dose (cigarettes per day) and duration of exposure (years as a smoker) in current smokers. Individuals who had smoked for a longer duration were more likely to be heavier smokers (age-and sex-adjusted linear regression Beta = 0.38 cigarettes per day for each year as a smoker P = 2.28 x 10^−8^). To minimise confounding between dose and duration of exposure, data for current smokers were split based on the median dose to generate time-specific subsets of heavy and light smokers at each exposure time point. The proportion of *smoker-enriched* cluster assignments increased with duration of exposure in both dose groups, stabilising at 15-19 years of exposure in heavy smokers, and 25-29 years in lighter smokers (**Supplementary Table 1**). The proportion of individuals assigned to the *smoker-enriched* cluster over time in heavy smokers was significantly greater than that in light smokers (Wilcoxon signed rank test P = 0.002; **Figure 2**).

### Clustering of former smokers depends on dose and time since cessation

Of the 1,466 former smokers assessed, 359 (24.4%) were assigned to the *smoker-enriched* cluster. The proportion of *smoker-enriched* cluster assignments decreased as time since smoking cessation increased (**Figure 3, Supplementary Table 2**). The highest proportion of *smoker-enriched* cluster assignments (64.4%) was observed in individuals who had quit smoking within a year prior to sampling. The proportion of *smoker-enriched* cluster assignments fell below 50% by 1 year following cessation. Contrary to the findings in current smokers, there was a significant negative relationship between dose and duration of exposure in former smokers (age-and sex-adjusted linear regression Beta = -0.18 cigarettes smoked per day for each year as a smoker P = 1.46 x10^−13^). Samples were next split on the median dose at each cessation time point to obtain a high-dose and low-dose group. The proportion of *smoker-enriched* cluster assignments was significantly lower in the low-dose group relative to the high-dose group (Wilcoxon signed rank test P = 1.5 x 10^−4^). The proportion of *smoker-enriched* cluster assignments in the low-dose group was consistently below 50% (**Figure 3**). The proportion of *smoker-enriched* cluster assignments for former smokers exposed to a high dose fell below 50% two years following cessation. From five years following smoking cessation, the proportion of *smoker-enriched* cluster assignments stabilised in high- and low-dose groups. Of the 902 individuals who had quit at least 5 years prior to sampling, 141 (15.6%) were assigned to the *smoker-enriched* cluster. As duration of exposure was not considered here, the analysis was repeated substituting years with cessation with pack years (years as a smoker x packs smoked per day), revealing a similar trend (**Supplementary Table 4, Supplementary Figure 1**). Using pack years as a metric, the proportion of *smoker-enriched* cluster assignments in the low-dose group stabilised from two years following smoking cessation, compared to five years following cessation in the high-dose group.

**Figure 3:**
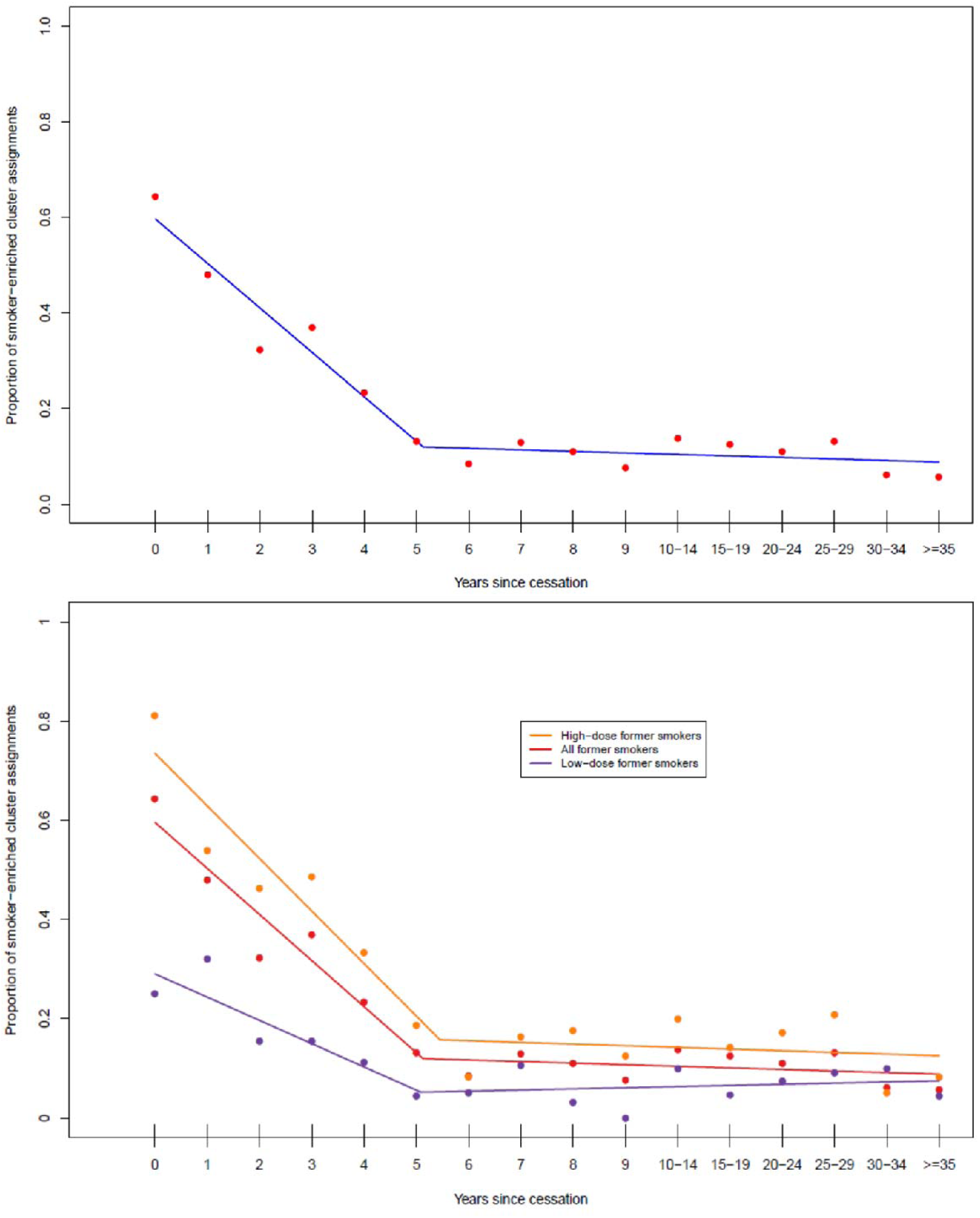
Proportion of former smokers assigned to Cluster 2 (smoker-enriched cluster) by years since smoking cessation. The upper panel displays a “broken stick” regression line for all samples. The lower panel displays “broken stick” regression lines for all samples (red), high-dose samples (orange) and low-dose samples (purple).

### Sensitivity analysis

To check the robustness of the predictions, two sensitivity analyses were considered (**Supplementary File 1**). In the first analysis, a parsimonious predictor was developed by selecting CpG sites that discriminated smokers from non-smokers with an AUC > 0.9 (five out of the 18,760 genome-wide EWAS sites identified by Joehanes et al. [5]). There was a slight improvement in the prediction of current versus never smokers using this score (**Supplementary Table 5**). However, the proportion of current smoker assignments in high-dose former smokers was consistently higher in the five-CpG predictor compared with the cluster-based predictor. Moreover, low-dose former smokers displayed a consistent proportion of current smoker assignments over time in comparison to the cluster-based assignments (**Supplementary Figure 3**).

In the second analysis, two predictors were developed based on polygenic scores for a subset of the most significant smoking-associated CpG sites (n = 90), and all smoking-associated sites (n = 17,529) [5]. The 90-probe polygenic predictor yielded similar results to the cluster-and AUC-based predictors (**Supplementary Table 6, Supplementary Figures 4-5**). In contrast, the polygenic score derived from the larger probe set displayed poorer predictions (**Supplementary Table 7, Supplementary Figures 6-7**).

## Discussion

In this study, we showed that smoking-based DNA methylation patterns are time- and dose-dependent. We identified two clusters from DNA methylation data in over 4,900 individuals – one enriched for current smokers and another enriched for never-smokers. It took 15-19 years for the majority of low-dose smokers to display a methylation profile that assigned them to the smoker-enriched cluster. It took less than 1 year for the majority of low-dose ex-smokers to be assigned to the never smoker-enriched cluster. By contrast, it took 5-9 years for the majority of heavy-dose smokers to display DNA methylation profiles corresponding to the smoker-enriched cluster, and up to 2 years since quitting before the majority of heavy-dose ex-smokers had methylation patterns that more strongly resembled those of never smokers. Furthermore, there is little impact of smoking dose on methylation-based clustering of smoking for those who had smoked for more than 25 years or for those who had stopped smoking for at least 7 years.

These findings suggest that a prolonged period of exposure to cigarette smoke is required before a smoking-related signature can be reliably identified using DNA methylation data. This is supported by evidence from multiple studies, which have reported an association between duration of exposure to cigarette smoke and an increased risk of oesophageal, lung and bladder cancers [22, 23, 24]. Moreover, a longer duration of exposure has been linked to an increased risk of chronic obstructive pulmonary disorder (COPD) and respiratory symptoms [25]. Should the DNA methylation profile of smokers be associated with an increased risk of smoking-related pathologies, the current findings suggest there is a dose-dependent period of exposure within which this risk is comparable to that of never smokers.

Others have reported reversion of smoking-associated DNA methylation changes in former smokers persisting beyond 30 years from cessation, with the most rapid reversion rates occurring in the within the first 14 years [26]. Moreover, increased methylation levels at *AHRR* has been reported in smokers undergoing cessation therapy [27]. Examination of cluster assignments in former smokers revealed to some degree the reversible nature of smoking-associated DNA methylation changes. Former light smokers were more likely to be assigned to the never smoker-enriched cluster, regardless of time since cessation. In contrast, a period of two years was required before the rate of never-smoker cluster assignments for former heavy smokers reached more than 50%.

A small proportion of never smokers were assigned to the smoker-enriched cluster. Such misclassifications may be a result of passive smoking, or other lifestyle-related correlates of smoking status. Although we did not observe an association between alcohol consumption and assignment of never smokers to the *smoker-enriched* cluster, it is possible that additional smoking-associated factors contribute to their misclassification. The effects of passive (i.e. second-hand) smoking on DNA methylation have been well established, with differential DNA methylation reported to persist up to decades following exposure to cigarette smoke in-utero [11, 12, 13]. It was not possible to determine whether the individuals profiled in the current study were exposed to cigarette smoke in utero as maternal smoking data were unavailable. Second-hand smoke exposure has also been linked to differential DNA methylation in adults. Similar DNA methylation patterns have been observed in lung tumours of smokers and second-hand smokers [28]. Hypomethylation of *AHRR* at cg05575921 has been linked to recent exposure to second-hand smoke [29], while others have reported significant associations between second-hand smoke exposure and differential DNA methylation in bladder cancer [30]. There was no association between misclassification of never smokers and co-habitation with, or other exposure to, smokers. However, information on the duration of co-habitation with smokers was not available, and there was no information regarding co-habitants and exposures prior to sampling.

We showed the use of AUC-based prediction of current/never smoking status using five probes is more accurate than the cluster-based prediction. However, smoking-associated DNA methylation changes at four of these five probes have been reported to persist decades following cessation [5]. Moreover, as the prediction thresholds for the five CpGs were selected to discriminate smoking status in the current sample, this generates a biased predictor when applied to the same data. While the predictive performance of the cluster-based predictions are less accurate for current/never smokers, its application to former smokers may be more suitable due to the inclusion of sites with reversible smoking-associated DNA methylation changes. This is reflected by the consistently higher proportion of current smoker assignments over time from the AUC-based predictor relative to the cluster-based predictor.

In a further sensitivity analysis, Z-score based polygenic methylation scores were built from all 18,760 genome-wide significant CpGs (n = 17,529 present in the GS:SHFS dataset) and also from the top 100 CpGs (n = 90 present in the GS:SHFS dataset). In the primary analysis, clusters were defined in relation to the methylation values in the Generation Scotland cohort, which may have introduced ascertainment bias. The polygenic analysis for the 90 CpGs yielded very similar results to the primary models. In contrast, the polygenic analysis derived from 17,529 probes did not perform as well. This was possibly due to the introduction of noise from many features of small effects. Conversely, predictive performance was improved by the inclusion of fewer features of larger effects.

In addition to the lack of information regarding maternal and past exposures to second-hand smoke, a further limitation to the current analysis is the presence of confounding between cigarette dose and duration of exposure to cigarette smoke. In order to minimise this association and to focus primarily on duration of exposure, the sample was stratified on the median dose at each time point assessed. A strength of this study is the use of a large and homogeneous analysis cohort. The Generation Scotland cohort comprises participants across a broad age range (18-99 years) which has permitted the analysis of smoking exposure in a large number of both recently-started and long-term smokers, as well as recently-quit and long-term former-smokers. Moreover, future analysis of smoking phenotypes and related health outcomes are possible, as a result of data linkage capabilities and sample collection for longitudinal DNA methylation profiling.

In conclusion, our findings suggest there is a dose-dependent interval within which smoking-associated DNA methylation are established. Furthermore, we have demonstrated a degree of reversibility of these changes in former smokers, whereby the interval of reversion is dependent on dose prior to smoking cessation. Consideration of duration of exposure in current smokers, and years since cessation in former smokers, coupled with dose, all measured via DNA methylation patterns, may assist in determining and stratifying risk of smoking-associated morbidities.

## Acknowledgements

This work was supported by Alzheimer’s Research UK Major Project Grant [ARUK-PG2017B-10]. Generation Scotland received core funding from the Chief Scientist Office of the Scottish Government Health Directorates [CZD/16/6] and the Scottish Funding Council [HR03006]. We are grateful to all the families who took part, the general practitioners and the Scottish School of Primary Care for their help in recruiting them, and the whole Generation Scotland team, which includes interviewers, computer and laboratory technicians, clerical workers, research scientists, volunteers, managers, receptionists, healthcare assistants and nurses. Genotyping of the GS:SFHS samples was carried out by the Genetics Core Laboratory at the Wellcome Trust Clinical Research Facility, Edinburgh, Scotland and was funded by the Medical Research Council UK and the Wellcome Trust (Wellcome Trust Strategic Award “STratifying Resilience and Depression Longitudinally” (STRADL) [104036/Z/14/Z]. DNA methylation data collection was funded by the Wellcome Trust Strategic Award [10436/Z/14/Z]. The research was conducted in The University of Edinburgh Centre for Cognitive Ageing and Cognitive Epidemiology (CCACE), part of the cross-council Lifelong Health and Wellbeing Initiative [MR/K026992/1]; funding from the Biotechnology and Biological Sciences Research Council (BBSRC) and Medical Research Council (MRC) is gratefully acknowledged. CCACE supports Ian Deary, with some additional support from Dementias Platform UK [MR/L015382/1]. AMM and HCW have received support from the Sackler Institute.

